# Transcriptional responses to arbuscular mycorrhizal symbiosis development are conserved in the early divergent *Marchantia paleacea*

**DOI:** 10.1101/2020.12.14.422721

**Authors:** Mara Sgroi, Uta Paszkowski

## Abstract

Arbuscular mycorrhizal symbiosis (AMS) arose in land plants more than 400 million years ago, perhaps acting as a major contributor to plant terrestrialization. The ability to engage in AMS is evolutionarily conserved across most clades of extant land plants, including early diverging bryophytes. Despite its broad taxonomic distribution, little is known about the molecular components that underpin AMS in early diverging land plants as the mechanisms regulating the symbiosis were primarily characterized in angiosperms. Several AMS associated genes were recently shown to be conserved in liverworts and hornworts, but evidence of them being associated with symbiosis in bryophytes is scarce. In this study, we characterised the dynamic response of the liverwort *Marchantia paleacea* to *Rhizophagus irregularis* colonization by time-resolved transcriptomics across progressive stages of symbiosis development. Gene orthology inference and comparative analysis of the *M. paleacea* transcriptional profile with a well characterised legume model -*Medicago truncatula* - revealed a deep conservation of transcriptional responses to AMS across distantly related species. We identified evolutionarily conserved patterns of expression of genes required for pre-symbiotic signalling, intracellular colonization and symbiotic nutrient exchange. Our study demonstrates that the genetic machinery regulating key aspects of symbiosis in plant hosts is largely conserved and coregulated across distantly related land plants. If bryophytes are confirmed to be monophyletic, our analysis provides novel insights on the first molecular pathways associated with symbiosis at the dawn of plant colonization of land.

**Significance Statement:** Arbuscular mycorrhizal symbiosis (AMS) between plants and soil fungi was proposed as one of the key adaptations enabling land colonization by plants. The symbiosis is widespread across most extant plant clades, including early-diverging bryophytes, suggesting that it evolved before the last common ancestor of land plants. Recent phylogenetic analyses uncovered that genes regulating AMS in angiosperms are present in the genomes of bryophytes. Our work shows that a set of these genes are transcriptionally induced during AMS in liverworts. Based on the conservation of their transcriptional profiles across land plants, we propose that these genes acquired an AMS-associated function before the last common ancestor of land plants.

## Introduction

When plants first colonized land between 515 and 470 mya, they overcame the challenge of surviving on a nutrient-deficient terrestrial substrate, with little to no organic matter available (1). How the first land colonizers managed to inhabit this new inhospitable environment is still unsure, but a growing body of evidence supports the hypothesis that plant-fungus symbioses were a pivotal component of this evolutionary transition (2–9). Identifying the molecular components that control extant plant-fungal symbioses might thus point towards the molecular mechanisms that made the first plant pioneers successful land colonizers. Arbuscular Mycorrhiza is the most widespread and well characterised form of plant-fungal symbiosis on Earth, engaging more than 80% of land plants across all major embryophyte clades (10). Fossil and phylogenetic data point to Arbuscular Mycorrhiza Symbiosis (AMS) being a monophyletic trait, which evolved between 450–407 million years ago in the last common ancestor of land plants (3, 7, 8, 11–14). The outcome of AMS is mutualistic: plants provide Glomeromycotina fungi with photosynthetically fixed carbon (carbohydrates and fatty acids) in exchange for mineral nutrients extracted from the rhizosphere (15–20).

Our current understanding of the molecular components that regulate AMS is based on evidence from a few angiosperm models, while molecular characterization of other tracheophytes and early-diverging bryophytes is at its dawn (14, 21–23).

AMS development includes three consecutive modules: 1. pre-symbiotic recognition, 2. tissue invasion and 3. nutrient exchange. During pre-symbiotic signalling both plant and fungal symbionts exude a broad range of signalling compounds in the rhizosphere to prime their symbiotic partner for symbiosis (24). Strigolactones are the most characterised component of plant exudates: their biosynthesis is upregulated under nitrogen or phosphate starvation and their perception triggers activation of fungal metabolism and hyphal branching (25–29). Upon perception of strigolactones, fungi boost the release of short-chain chitin oligosaccharides and small secreted proteins that act as signalling molecules for plant perception of AMF (30, 31). The downstream signalling cascade activated upon fungal perception is shared by both AMS and root nodule symbiosis and takes thus the name of Common Symbiosis Signalling Pathway (CSSP). Through a complex nuclear signalling cascade, the CSSP culminates in the interaction of MtIPD3/LjCYCLOPS with DELLA transcription factors to activate transcriptional reprogramming of the host cell (reviewed in (32)). DELLA transcription factors were first discovered for their role in repressing Gibberellic Acid (GA) signalling, and both exogenous addition of GA to AMS roots and genetic mutation of *DELLA* genes lead to impaired arbuscule development, suggesting that GA is involved in regulating AMS through its interaction with DELLA proteins (33–36).

CYCLOPS/DELLA work in concert to induce a plethora of GRAS transcription factors required to regulate strigolactone biosynthesis, lipid biosynthesis, arbuscule formation, and symbiotic nutrient transport (37–39). The gene list includes: *Reduced Arbuscular Mycorrhiza 1* (*RAM1*), *Reduced Arbuscule Development 1* (*RAD1*), *Nodulation Signalling Pathway 1* (*NSP1*), and *NSP2* (reviewed in 39).

CSSP activation shifts the plant into a permissive state, in which the fungus is able to intracellularly colonize host tissues and form highly branches hyphal structures, “arbuscules”, that specialise in nutrient exchange. Most of the symbiosis-dependent nutrient exchange occurs across the peri-arbuscular membrane (PAM), which is highly enriched with several transmembrane transporters for phosphate (MtPT4/OsPT11), ammonium uptake (MtAMT2;3/ SbAMT3;1), and lipid (MtSTR, MtSTR2) and glucose (MtSWEET1b) efflux (40–46). All known symbiosis-specific nutrient transporters are transcriptionally induced in AMS colonized tissue, but the dynamics of nutrient exchange seem to be further regulated at the PAM interface. Indeed, mutations in the PAM-specific *Arbuscular Receptor-like Kinase 1* (*OsARK1/MtKIN3*) significantly reduce vesicle formation and overall fungal colonization in rice and *M. truncatula*, suggesting that signalling at the PAM is necessary to maintain fungal fitness (22, 47).

If AMS is a basal trait of land plants, the underpinning molecular pathways should be conserved across extant AMS-competent clades as a result of positive selection, while they should be absent in plant lineages that no-longer associate with AMS as a result of co-elimination (14, 48). Three independent phylogenomic studies demonstrated this hypothesis by discovering that a core set of genes are consistently retained across AMS-competent angiosperms but lost in AMS-incompetent lineages (22, 49, 50). The latest and most stringent of these phylogenomic analyses identified 72 orthogroups conserved across AMS-competent angiosperms (22). Several gene families identified by the study have a characterized molecular function in AMS, whilst others display a reduced colonization phenotype in *M. truncatula* mutants but their function has not yet been characterized (22). The majority (86%) of *M. truncatula* genes belonging to these orthogroups are transcriptionally upregulated in *M. truncatula* AMS, suggesting that evolutionary conservation in AMS lineages coupled with transcriptional induction are reliable predictors to identify genes required for symbiosis (22).

Recent large-scale phylogenetic analyses went beyond the flowering plants clade to identify symbiosis genes conserved across all land plants engaging in AMS (14, 21). A high degree of sequence conservation was observed within land plants: orthologs of the major components of the CSSP (*LysM-RLKs, SYMRK/DMI2, DMI1, CcaMK* and *CYCLOPS/IPD3*) and most GRAS transcription factors (*RAD1, RAM1, NSP1, NSP2*) are conserved in AMS-competent bryophytes (12, 14, 21). Bryophyte orthologs of several PAM-associated proteins and a lipid biosynthesis gene (*VAPYRIN, LIN, SYNTAXIN, STR* and *STR2, RAM2*) were also identified in these studies, suggesting that the molecular components necessary for intracellular accommodation and nourishment of fungal symbionts evolved before the last common ancestor (LCA) of land plants (14, 21). A small subset of these evolutionarily conserved genes (*SymRK, CCaMK, CYCLOPS, RAD1, STR and STR2*) also follow the same co-elimination trend that characterises their angiosperm orthologs, as they are consistently lost in non-host embryophyte clades (14). The finding that some of these evolutionarily conserved genes (*LcNSP1*, *LcRAD1*, *LcRAM2*, *LcSTR*, and *LcSTR2*) are also upregulated during AMS in the liverwort *Lunularia cruciata* hints to a certain degree of conservation of AMS transcriptional responses in bryophytes (21).

Taken together, sequence conservation in bryophytes suggests that the genes required for angiosperm AMS evolved before the LCA of land plants. It is however still to be determined whether the majority of genes conserved in early-diverging plants associate with bryophyte AMS, or if instead they fulfil distinct cellular functions and were co-opted for symbiosis after the divergence of the bryophyte clade.

Based on the observation that most genes conserved for symbiosis across angiosperms are upregulated during *M. truncatula* AMS (22), we investigated AMS transcriptional responses in bryophytes using *M. paleacea* as a liverwort model. Through an RNAseq time-course we characterised symbiosis development in *M. paleacea* and compared gene expression profiles to corresponding *M. truncatula* orthologs. Through this comparative approach we identified a core set of AMS genes that are not only conserved in liverworts but also transcriptionally induced in response to AMS. The conservation of this core symbiotic gene set in bryophytes suggests that it evolved before the LCA of embryophytes, providing novel insights into the molecular toolkit that was co-opted for symbiosis at the origin of land plants.

## Results

### Transcriptional responses to AMS in M. paleacea intensify over time

To investigate the transcriptional response of *M. paleacea* during AMS, we performed RNAseq of colonized thalli at 5-, 8- or 11-weeks post inoculation (WPI) with *R. irregularis*. For every time point, we compared transcript levels of genes of mock-inoculated thalli to colonised *M. paleacea* thalli. The predominant fungal structure observed in colonised thalli at 5WPI was intracellular hyphae, with arbuscule levels increasing in abundance at 8WPI (Fig. 1a). By 11WPI the midrib area of *M. paleacea* thalli was intensely colonised, with abundance of all quantified fungal structures: hyphae, arbuscules and vesicles (Fig. 1a). A distinctive red pigmentation specific to colonized *M. paleacea* thalli (51), accumulated proportionally to intracellular colonization levels (Fig. 1a).

**Figure 1.**
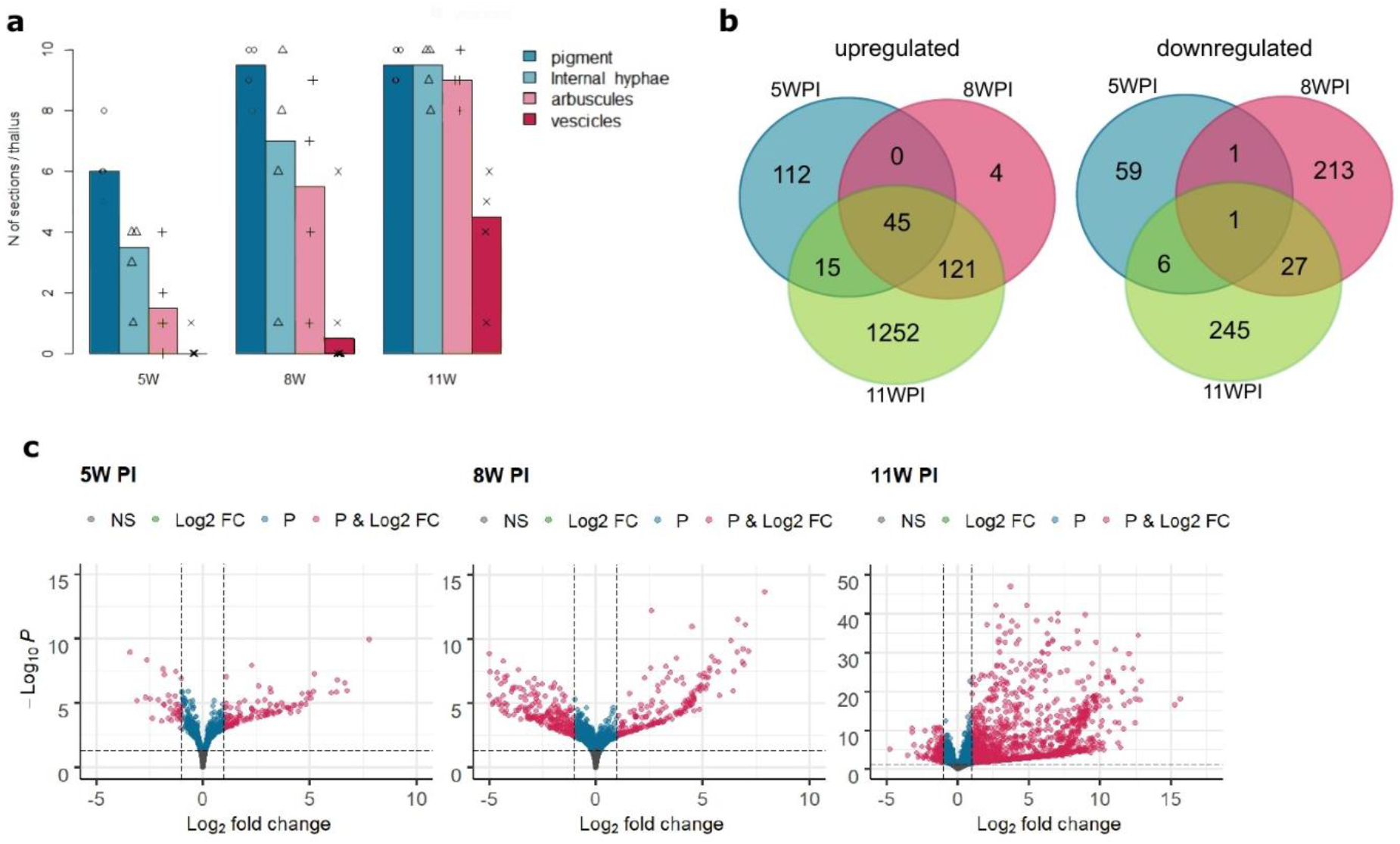
Transcriptional responses of *M. paleacea* to *R. irregularis* colonization. **a.** Arbuscular mycorrhiza colonisation levels of WT *M. paleacea* thalli inoculated with *R. irregularis* at 5, 8 and 11 weeks post-inoculation (WPI). Each datapoint represents the number of sections per thallus in which the fungal structure was observed. Each independent replicate was screened for all fungal structures over a total of 10 sections. Bars represent the average number of observations of each fungal structure across all replicates; **b.** Venn diagrams of differentially expressed genes (adjusted p-value < 0.05, Log2 fold change > |1|) in pairwise comparisons of mock and *R. irregularis-*colonised thalli at 5WPI, 8WPI and 11WPI; **c.** Volcano plots of differentially expressed genes in pairwise comparisons of mock and *R. irregularis-*colonised thalli at 5WPI, 8WPI and 11WPI. Significantly differentially expressed genes are displayed in magenta. NS = non-significant; Log2 FC = past the Log2 fold change threshold; P = past the P-value threshold; P & Log2 FC = past Log2 fold change and adjusted P-value threshold.

Differential expression analysis revealed that the transcripts of a relatively small pool of genes are significantly changing in abundance (adjusted p-value<0.05 & Log2 fold change >|1|) between control and colonised plants at early (5WPI) and intermediate (8WPI) stages of symbiosis (Fig.1 b,c). At 5WPI, the proportion of genes significantly upregulated in response to fungal colonisation (172 genes) was twice the volume of the downregulated pool (67 genes), suggesting that an AM-specific transcriptional response is initiated at early stages of symbiosis. The transcriptional response at 5WPI can be divided into two core clusters: one composed of genes specific to early symbiosis (112 genes) and one smaller group of genes (45 genes) consistently upregulated across all stages of AMS, independent from changes in intracellular fungal structures over time (Fig.1b). A set of 121 genes are significantly upregulated at intermediate and late stages of symbiosis (Fig.1b). As these later time-points are characterised by a greater abundance of arbuscules, genes associated with arbuscule development and nutrient exchange are likely to be overrepresented in this cluster (Fig. 1a, b).The strongest response to fungal colonisation was observed at 11WPI, which displays a distinctive bias towards upregulated genes compared to downregulated genes (Fig. 1b, c). This pattern of gene expression is characteristic of AMS and tends to be associated with the expression of AMS-specific transcripts only induced in mycorrhizal conditions (52–54).

### GO enrichment analysis reveals evolutionarily conserved pathways associated with AMS

To investigate the identity of transcripts differentially expressed (DE) at each stage of AMS, the transcriptome of *M. paleacea* was functionally annotated with Trinotate (55). The list of significantly upregulated genes specific to early symbiosis comprises a high proportion of candidate pathogenesis-related (PR) proteins, including peroxidases, protease inhibitors and chitinases (Supplementary table 1). Several PR protein families induced in *Marchantia polymorpha* by oomycete infection (56) were also significantly induced at early stages of mycorrhizal symbiosis in *M. paleacea*: PR6a (*MPA27867*, FC = 338.50), PR9 (*MPA22356*, FC= 12.54), PR2 (*MPA28496*, FC = 7.53), PR5 (*MPA25727*, FC = 5.54) suggesting a degree of overlap between defence and symbiosis signalling (Supplementary tables 1,7). These PR proteins were significantly downregulated at 8WPI, indicating that initial defence responses were actively dampened over time (Supplementary table 1). Amongst the genes significantly upregulated at 5WPI were also a number of genes encoding protein families involved in regulation of plant defence genes: two serine peptidase inhibitors (SERPINs) (*MPA17316*, FC= 3.86; *MPA17337*, FC= 2.70) and three E3 ubiquitin ligases (*MPA12322*, FC = 10.56; *MPA5979*, FC = 4.30; *MPA15840*, FC = 4.01).

Several serine/threonine kinases were significantly upregulated at 5WPI (*MPA17978*, FC= 7.16; *MPA19735*, FC= 3.696; *MPA16100*, FC= 2.690; *MPA18092*, FC= 2.60; *MPA7445*, FC = 2.507; *MPA8794*, FC = 2.059; *MPA6760*, FC = 2.035) indicating an induction of signalling in response to intracellular colonization by *R. irregularis*. Intracellular signalling might be coupled with transcriptional reprogramming as two DNA-binding TFs (*MPA13879*, FC = 3.14; *MPA25954*, FC = 3.11) are also significantly upregulated at 5WPI (Supplementary table 1). A number of genes induced at 5WPI are annotated as components of the phenylpropanoid pathway, including a phenylalanine ammonia lyase (*MPA10826*, FC = 10.93), a naringenin-chalcone synthase (*MPA18004*, FC = 45.19), and a trans-cinnamate 4-monooxygenase (*MPA12664*, FC = 2.19). The upregulation of genes involved in the phenylpropanoid pathway coincided with accumulation of the cell wall pigmentation characteristic of colonized *M. paleacea* thalli (Fig. 1a)(51), suggesting that the pigment might be of phenylpropanoid origin.

We performed GO Enrichment analysis to identify molecular functions and biological processes upregulated during *M. paleacea* AMS (Supplementary table 2). Analysis of the list of genes significantly upregulated across all stages of AMS identified “gibberellin biosynthetic process” as the most overrepresented GO annotation in the dataset (FDR = 2.72E-06), closely followed by “nutrient reservoir activity” (FDR = 4.90E-04) (Supplementary table 2). These findings suggest that nutrient exchange and storage are transcriptionally regulated throughout the symbiosis. Analysis of the time-point with the highest transcriptional response to symbiosis (11WPI) highlighted two additional overrepresented membrane-specific terms: “proton export across plasma membrane” (FDR = 0.023) and “proton-exporting ATPase activity” (FDR = 0.027), in agreement with the high abundance of arbuscules observed at this late AMS stage (Fig. 1a)(Supplementary table 2). The GO terms “peptidase activity” (FDR = 0.019) and “aspartic-type endopeptidase activity” (FDR =0.015) were significantly enriched at this time point, in accordance with observations that several AM-induced proteases accumulate in or adjacent to arbusculated cells in vascular plants (57–60). The term “nucleosomal DNA binding” is also significantly enriched at this late stage of AMS (FDR = 0.03), indicating that fungal colonization might be linked to chromatin remodelling (Supplementary table 2).

Taken together, analysis of GO annotation in *M. paleacea* AMS suggests that a transient induction of pathogenesis-related proteins, serine/threonine kinase-mediated signalling, gibberellin-like phytohormone metabolism and chromatin remodelling could be a shared response of liverworts and vascular plants to AMS.

### Orthology inference and comparative analysis reveal deep homologies in transcriptional responses to AMS

Since GO enrichment analysis suggested consistent similarities in the transcriptional profiles of *M. paleacea* and vascular plants, we performed gene orthology inference to directly compare expression patterns of AMS genes across species. We selected 16 genomes representative of major land plant clades, maintaining a balanced representation of AMS host and AMS non-host clades, with two Charophyte algae as an outgroup (Supplementary Fig. 1, Supplementary Table 3). To increase phylogenetic support of bryophyte clades we included four bryophyte transcriptomes from the 1,000 plants (1KP) project (61)(Supplementary Fig. 1, Supplementary Table 3).

Our Orthofinder pipeline identified a total of 10057 orthogroups containing *M. paleacea* genes (Supplementary table 4). For the purpose of this report, we focused our analysis on the comparison between *M. paleacea* and the dicot AMS model *M. truncatula. M. paleacea* shares overall 6340 orthogroups with *M. truncatula*, comprising 1265 single-copy orthologs (SCOs)(Supplementary table 5). We compared expression patterns of SCOs by plotting the log2 fold change (FC) of *M. paleacea* DE genes in late symbiosis (11WPI) against the log2 FC of DE genes in *M. truncatula* roots at 27 days post inoculation (DPI) with *R. irregularis* (19)(Fig. 2a). The 27 DPI time point from Luginbuehl *et al.* (2017) was selected for this meta-analysis as it displays the highest abundance of all fungal structures (intracellular hyphae, arbuscule, vesicle) observed in this study at 11WPI (Fig. 1a).

**Figure 2.**
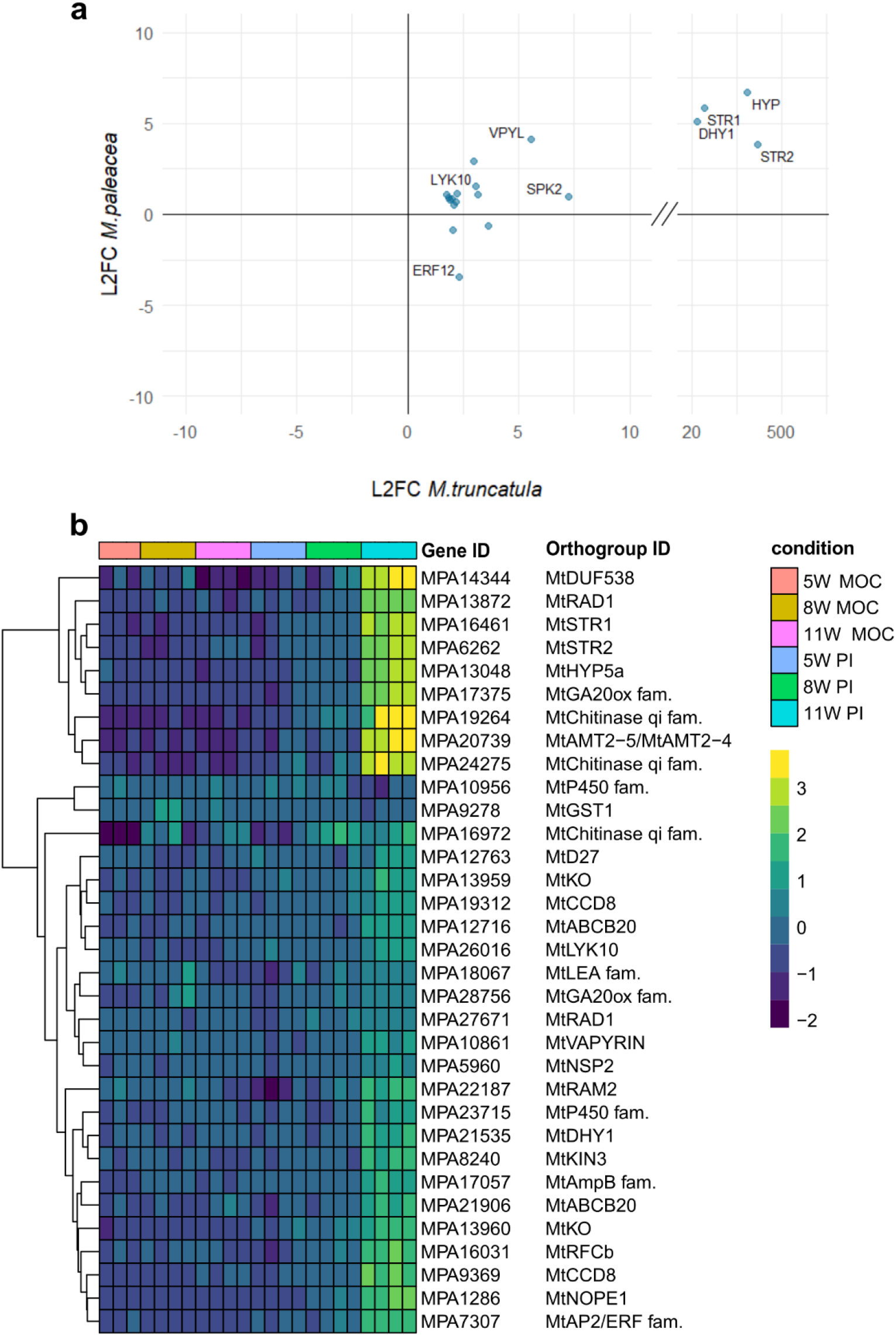
Comparison of AMS transcriptional responses in *M. paleacea* and *M. truncatula* colonized by *R. irregularis.* **a.** Scatterplot displaying the direction of Log2 fold change (L2FC) in gene expression of *M. paleacea* and *M. truncatula* single-copy orthologs differentially expressed during AMS colonization. *M. truncatula* L2FC values obtained from Luginbuehl *et al*. (2017). Only orthologs significantly differentially expressed in both species (L2FC>|1|, adjusted P-value < 0.05) are shown; **b.** Heatmap of differentially expressed *M. paleacea* genes with a *M. truncatula* ortholog upregulated during AMS colonization. Only sequences with known gene names in *M. truncatula* are displayed for ease of representation, full figure available in Supplementary Fig. 2.

Our comparison identified 12 SCOs that were significantly DE (P-adj <0.05 & Log2 FC >|1|) in *M. paleacea* and *M. truncatula* during AMS (Fig. 2a, Table 1). Eleven genes were upregulated in both species, and half of them (6/11) encode for proteins involved in fatty acid biosynthesis or transport, including the two symbiosis-specific half ABC transporters STR/STR2 (*MPA16461*, FC = 56.53; *MPA6262*, FC = 14.42) and the short-chain dehydrogenase/reductase DHY1 (*MPA21535*, FC = 34.46)(Fig. 2a, Table 1). The LysM-domain receptor kinase LYK10 is conserved and upregulated in *M. paleacea* (*MPA26016*, FC = 2.880), suggesting that LysM receptor kinases might be involved in regulating symbiosis in bryophytes. Finally, both orthologs of the PAM-associated protein VAPYRIN are co-upregulated in *M. paleacea* (*MPA10861*, FC = 17.38) and *M. truncatula* AMS (Fig. 2a, Table 1). Taken together, this SCO analysis suggested that several aspects of AM symbiosis are conserved across land plants. However, this approach is limited by the evolutionary distance between *M. paleacea* and *M. truncatula* and by the occurrence of multiple duplication events in tracheophytes. Indeed, only a small proportion of *M. paleacea* genes correspond to a single ortholog in *M. truncatula* (13.42%) whilst the majority of shared orthogroups display more complex or one-to-many (20.26%) or many-to-many (66.33%) relationships between orthologous sequences (Supplementary Table 5).

**Table 1.** List of Single copy orthologs differentially expressed in *M. paleacea* and *M. truncatula* during AMS. Only genes significantly DE (adjusted p-value<0.05 & Log2 fold change>|1|) in both *M. paleacea* and *M. truncatula* (19) are shown; L2FC = Log2 fold change; 11WPI = 11 weeks post inoculation; 27DPI= 27 days post inoculation.

| *M. paleacea*<br>gene ID | *M. paleacea*<br>(11WPI) L2FC | *M. truncatula*<br>gene ID | *M. truncatula*<br>(27DPI) L2FC<sup>+</sup> | *M. truncatula* gene<br>annotation | *M. truncatula*<br>Gene name |
| --- | --- | --- | --- | --- | --- |
| MPA6262 | 3.850 | Medtr5g030910 | 364.363 | white-brown-complex ABC transporter family* | STR2 |
| MPA27420 | 6.722 | Medtr4g035250 | 306.007 | hypothetical protein | HYP |
| MPA16461 | 5.821 | Medtr8g107450 | 70.274 | white-brown-complex ABC transporter family* | STR1 |
| MPA21535 | 5.107 | Medtr4g097510 | 28.68 | enoyl-(acyl carrier) reductase* | DHY1 |
| MPA18294 | 0.998 | Medtr8g074920 | 7.263 | MtSPK2 serine/threonine receptor like kinase | SPK2 |
| MPA10861 | 4.119 | Medtr6g027840 | 5.570 | ankyrin repeat RF-like protein, putative | VPY |
| MPA24714 | 1.069 | Medtr3g076630 | 3.157 | pyruvate dehydrogenase E1 beta subunit* | PDHB |
| MPA26016 | 1.526 | Medtr5g033490 | 3.097 | LysM type receptor kinase | LYK10 |
| MPA26315 | 2.936 | Medtr1g105965 | 2.991 | pyruvate kinase family protein* |  |
| MPA24462 | -3.459 | Medtr2g014300 | 2.310 | AP2/ERF TF ethylene response factor | ERF12 |
| MPA16240 | 1.173 | Medtr8g024310 | 2.229 | pyruvate dehydrogenase E1 alpha subunit* | PDHA |
| MPA19145 | 1.066 | Medtr7g078070 | 1.772 | cysteine synthase/L-3-cyanoalanine synthase |  |

\* genes involved in fatty acid biosynthesis/transport; <sup>+</sup> *M. truncatula* L2FC data from Luginbuehl *et al.* (2017).

To better understand the conservation of AMS transcriptional profiles independently of clade-specific duplications, we filtered our dataset to display only genes significantly DE in *M. paleacea* with at least one *M. truncatula* ortholog DE in the RNAseq dataset of Luginbuehl *et al.* (19). This analysis identified 178 *M. paleacea* genes with at least one ortholog upregulated in *M. truncatula* AMS (Supplementary fig. 2, supplementary table 6). The analysis revealed that the *M. paleacea* orthologs of the core strigolactone biosynthetic genes *D27* (*MPA12763*, FC = 2.76), *CCD8a* (*MPA9369*, FC = 6.70), *CCD8b* (*MPA19312*, FC = 2.46), as well as the GlcNAC transporter *NOPE1* (*MPA1286*, FC = 476.06) are conserved and significantly upregulated in *M. paleacea* during AMS (Fig. 2b, Supplementary table 6), suggesting that the requirement of these genes for plant rhizosphere signalling is ancestral.

Orthologs of the two GRAS transcription factors *RAD1* (*MPA13872*, FC = 34.30) and *NSP2* (*MPA5960*, FC = 2.34) were co-upregulated in *M. paleacea* and *M. truncatula* datasets, suggesting they are required for symbiosis in bryophytes as well as in angiosperms (Fig. 2b, Supplementary tables 5,6).

Our analysis identified one transcription factor from the AP2/ERF2a family (*MPA7307*) that, like *RAD1a*, is exclusively induced in colonized tissue and significantly upregulated at late stages of symbiosis in both *M. paleacea* (FC = 424.61) and *M. truncatula* (Fig. 2b, Supplementary table 6). Since the three *M. truncatula* orthologs in this AP2/ERF2a clade (*Medtr4g082345*, *Medtr6g012970*, *Medtr7g011630*) are only conserved in AMS-competent angiosperms (22), we investigated the evolutionary conservation of *MPA7307* in the non-host sister taxon of *M. paleacea*: *M. polymorpha*. We observed, with low phylogenetic support, that the *M. polymorpha* ortholog of *MPA7307* is missing from the *M. polymorpha* genome, whilst its closest paralog (*MPA21338*) is conserved in *M. polymorpha* (Supplementary fig. 3).

We finally investigated the molecular pathways known to regulate bidirectional nutrient exchange in angiosperms. We identified a conserved ortholog of *M. truncatula*’s *AMT2;4/AMT2;5* ammonia transporters, which is one of the most upregulated genes at 11WPI (*MPA20739*, FC = 1078.64). Furthermore, the *M. paleacea* orthologs of the serine/threonine kinase *MtKIN3*/*OsARK1* are significantly upregulated in response to AMS in *M. paleacea* (*MPA8240*, FC = 10.05) and *M. truncatula* (Fig. 2b, Supplementary tables 5,6). In addition to the lipid transporters and biosynthesis genes identified in the SCO comparison (Fig. 2a, Table 1), we further identified the *M. paleacea* ortholog of the fatty acid biosynthetic gene *RAM2*, which is significantly upregulated at late stages of symbiosis (*MPA22187*, FC = 4.36)(Fig. 2b, Supplementary tables 5,6). The upregulation of several fatty acid biosynthetic genes (Fig. 2b, Supplementary table 6) co-occurring with the accumulation of fungal vesicles (Fig.1a) suggests that host-derived lipids might be a conserved currency of AMS across land plants.

The *M. paleacea* orthologs of several *M. truncatula* genes predicted to have an active role in AMS due to their transcriptional profile, mutant phenotype and evolutionary conservation, are significantly DE at 11WPI in our dataset: *Replication Factor C* (*MPA16031*, FC = 12.85), *ABCB20 (MPA17057*, FC =12.60; *MPA12716*, FC =3.17), two cytochrome P450 family proteins *(MPA10956*, FC =0.28; MPA23715, FC = 3.56), three Chitinases *(MPA19264*, FC = 1131.48; *MPA24275*, FC = 225.97; *MPA16972*, FC = 3.40), *Hypothetical Protein 5A* (*MPA13048*, FC = 83.63) and *DUF538 (MPA14344*, FC = 58.89)(Fig. 2b, Supplementary tables 5,6)(22).

Despite the extensive evidence for conservation of transcriptional responses to AMS, we identified some differences between *M. paleacea* and *M. truncatula* AMS profiles that might reflect separate evolutionary trajectories in AMS gene families. No direct orthologs of the mycorrhiza-specific phosphate transporter MtPT4/OsPT11 were identified in *M. paleacea*. As AMS symbiosis was previously reported to increase the phosphorus uptake in *M. paleacea* thalli (51), we probed our RNAseq dataset to identify a different group of phosphate transporters induced during AMS. We detected two *M. paleacea* genes (*MPA20295*, *MPA19863*) belonging to the same orthogroup as the angiosperm PHT transporters, which were strongly and exclusively mycorrhiza induced (FC > 675)(Supplementary tables 1,4). We additionally identified two phosphate transporters upregulated at 11WPI that also showed a baseline expression level in mock (*MPA6710*, FC = 6.44; *MPA15906*, FC = 12.88). *MPA6710* and *MPA15906* belong to an orthogroup (OG0001191) which does not contain any angiosperm sequences aside from *A. trichopoda*, suggesting that no close ortholog of this clade is retained in other angiosperms (Supplementary table 4).

In a related observation, the *M. paleacea* single ortholog of MtHA1/OsHA1 identified by our analysis was not significantly DE during AMS (*MPA20871*, P-adj > 0.05)(Supplementary tables 1, 5). As MtHA1 activity is necessary to enable MtPT4’s function in *M. truncatula* (62), we investigated expression levels of other *M. paleacea* proton pumps. We identified three different HA proton pumps (*MPA25345, MPA28647 and MPA9460*) DE in our dataset, which belong to the same orthogroup as MpaHA1 but are more distantly related to MtHA1/OsHA1 (Supplementary tables 1,4,5). *MPA25345* and *MPA28647* were significantly induced from the earliest time-point (respectively FC = 106.45, FC = 10.81) and their expression level consistently increased with plant colonization levels (Supplementary table 1).

Taken together, this work uncovered a deep homology in the transcriptional response of liverworts and angiosperms to AMS. Our findings suggest that several molecular modules necessary for symbiosis establishment and maintenance in angiosperms - strigolactone biosynthesis, nuclear signalling, arbuscule maintenance and symbiotic nutrient exchange - are not only conserved but significantly induced in liverworts in response to AMS.

### Gibberellin-like diterpenoids might be induced during *M.paleacea* AMS

GO term enrichment analysis of the gene set upregulated across all stages of symbiosis highlighted an overrepresentation of terms associated with gibberellin biosynthesis (Supplementary table 2). Seven out of 13 genes annotated as gibberellin biosynthetic genes also presented the GO annotation “Gibberellin-20-oxidase” (GA20ox). Bryophytes have been previously reported to possess genes involved in GA precursor biosynthesis (*ent-copalyl diphosphate synthase* (*CPS*), *ent-kaurene synthase* (*KS*), *ent-kaurene oxidase* (*KO*), *ent-kaurenoic acid oxidase* (*KAO*)), while there is contrasting evidence on the existence of GA20ox, GA2ox and GA3ox orthologs and canonical GA bioactive molecules (63–71). We therefore investigated the identity of genes annotated as “Gibberellin-20-oxidase” and the expression levels of GA-precursor biosynthetic genes in *M. paleacea*. Orthology inference indicates that the *M. paleacea* transcripts that were mis-annotated by Trinotate as GA20ox are not orthologous to any other embryophyte sequence in our dataset, suggesting that they are not involved in an evolutionarily conserved biosynthetic pathway. Analysis of the *M. truncatula* GA20ox orthogroup instead highlighted a different set of bryophyte sequences at the base of the tracheophyte GA20ox clade, including three candidate *MpaGA20ox-like* orthologs (*MPA17375*, *MPA8132*, *MPA28756*) (Supplementary tables 4,5). We additionally identified all *M. paleacea* orthologs of GA-precursor biosynthetic genes based on the annotation of its sister taxon *M. polymorpha*: MpaCPS (*MPA24439*), MpaKS (*MPA15116*), two MpaKOs (*MPA13960*, *MPA29074*), and MpaKAO (*MPA18360*) (Supplementary table 7). DE analysis of these potential GA biosynthetic genes revealed that *MpaKOa* (*MPA13960*) was significantly upregulated across all stages of symbiosis (5W-FC = 3.84, 8W-FC = 6.25, 11W-FC = 16.81) whilst *MpaKOb* (*MPA29074*) was significantly DE at 11WPI (FC = 3.12). Two of the three candidate *MpaGA20ox* genes were also induced at 11WPI (*MPA17375*, FC = 81.91; *MPA28756*, FC = 2.29), whilst genes involved in earlier stages of GA biosynthesis – *MpaCPS* & *MpaKS* - were not induced (P-adj > 0.05) (Supplementary table 6). Taken together, evidence of the conservation and transcriptional induction of GA biosynthetic genes in *M. paleacea* suggests that some gibberellin-like diterpenes might be synthesised during AMS in liverworts.

## Discussion

This study discovered that transcriptional responses to AMS are deeply conserved across early-diverging land plants and angiosperms. By comparing the transcriptional profile of the liverwort *M. paleacea* to the profile of the AMS model *M. truncatula* we found that the overlap in molecular responses of land plants to AMS starts from pre-symbiotic signalling. Previous studies revealed that whilst strigolactone biosynthetic genes are conserved across land plants, the signalling components involved in plant perception of the phytohormone likely evolved independently in mosses and angiosperms (23, 72). Our finding that all strigolactone biosynthetic genes are upregulated during *M. paleacea* AMS supports the hypothesis that strigolactones first evolved as a rhizosphere signalling molecule in the LCA of land plants and were later co-opted as phytohormones (23, 72). We were also able to observe that the N-acetylglucosamine transporter *NOPE1* is evolutionarily conserved in *M. paleacea* and considerably upregulated during both *M. paleacea* and *M. truncatula* AMS, suggesting that the yet uncharacterised mechanism by which NOPE1 influences rhizosphere signalling is ancestral to land plants.

Downstream of reciprocal sensing, fungal hyphae establish contact and successfully penetrate host plant tissues. In our RNAseq dataset, intracellular colonization by *R. irregularis* significantly induces several *M. paleacea* PR protein-encoding genes that are also induced in response to *Phytophthora palmivora* infection in *M. polymorpha* (56). These findings suggest that the initial response of *M. paleacea* to fungal colonization partially overlaps with defence responses. Similar patterns of PR-protein induction were observed in a variety of angiosperm model species (73–79) and were linked to increased resistance of AMS hosts to pathogens (77, 79–82). In this study we demonstrate that the transcriptional activation of genes encoding PR protein is a conserved signature of bryophyte AMS, but whether AMS contributes to improved pathogen resistance in liverworts remains to be investigated.

In angiosperms, a regulatory network of transcription factors translates CSSP signalling into cellular reprogramming to accommodate fungal colonization (reviewed in (39)). Our study confirms that two GRAS TFs (*NSP2* and *RAD1*) are conserved in bryophytes and significantly induced during *M. paleacea* AMS, suggesting that GRAS family proteins play an active role in the regulation of symbiosis in bryophytes as well as in angiosperms. Two independent phylogenetic studies had previously identified single orthologs of GRAS TFs in liverworts (*NSP1, NSP2*, and *RAD1*) with Delaux *et al.* (2015) additionally claiming to have identified an ortholog of *RAM1* in the liverwort *L. cruciata* (21, 83). Our study identified orthologs of *RAD1, NSP1* and *NSP2* but no ortholog of *RAM1*, in agreement with the findings from Grosche *et al.* (83). In contrast to *NSP1/NSP2* expression levels in *L. cruciata* (21), *MpaNSP1* was not differentially expressed in our dataset, whilst *MpaNSP2* was significantly induced in AMS tissue, suggesting clade-specific differences in regulation of AMS genes. As *NSP2* transcription is regulated by the nutritional status of the plant in angiosperms (84, 85), differences in nutrient conditions between independent experimental setups might explain the observed discrepancies between *M. paleacea* and *L. cruciata*. Our analysis identified a novel transcription factor from the AP2/ERF2a family that, like *RAD1a*, is exclusively induced in colonized tissue and significantly upregulated at late stages of symbiosis (*MPA7307*). We observed that the *M. polymorpha* ortholog of *MPA7307* is missing from the *M. polymorpha* genome, whilst its closest paralog (*MPA21338*) is conserved in M. *polymorpha*. As the three Medicago orthologs of this AP2/ERF2a clade are amongst those genes specific to AMS-competent angiosperms (22) we suggest that the pattern of trait loss and co-elimination observed for this gene family might be conserved in liverworts.

Our analysis suggests that not only the transcriptional regulators of symbiosis are conserved and induced in *M. paleacea*, but also their downstream pathways. We confirmed the findings that the lipid biosynthesis and transport genes *RAM2*, *STR* and *STR2* and are conserved and upregulated during liverwort AMS (21). Our study additionally identifies a novel ammonium transporter orthologous to *MtrAMT2;4/MtrAMT2;5,* which is strongly induced in colonized tissues of *M. paleacea,* supporting the hypothesis that symbiotic nitrogen uptake is an evolutionarily conserved aspect of AMS (44, 86–90).

Whilst lipid and nitrogen transfer show evolutionarily conserved patterns of gene expression, a lower degree of homology was observed for phosphate transport. The AMS-inducible phosphate transporter and the three transmembrane proton pumps upregulated in *M. paleacea* are not orthologous to the mycorrhiza-specific transporters required for symbiosis development in angiosperms (*MtPT4/OsPT11, HA1*). Instead, the genes identified in this study are orthologs of the LcPT and LcHA transporters upregulated in the transcriptome of *L. cruciata* during AMS (21). These findings support the hypothesis that whilst most genetic and transcriptional responses to AMS evolved early in the history of land plants, clade specific recruitment of AMS genes is possible, and likely required to fine-tune symbiotic interactions to the physiological requirements of the host (21).

Gibberellin biosynthesis and signalling might be an example of such clade specific adaptations. We observed that a considerable proportion of annotated DE genes in *M. paleacea* mentioned GA biosynthesis, in conflict with the notion that GA active compounds are not known to be produced in bryophytes (64, 68–70). Amongst the significantly upregulated genes in our dataset we identified two orthologs of *KO* (required for synthesis of the GA-precursor *ent-*kaurene) and two orthologs of GA-20-oxydases (required to convert inactive GA12 precursors into bioactive GAs (70)). Similarly, the *M. truncatula* orthologs of *KO*, *KAO* and multiple GA-20-oxydases are upregulated during AMS (19). Whilst *KOs* were widely identified in bryophytes, there is contrasting literature on the existence of *sensu stricto* GA-20-oxidases (23, 64, 68, 69, 71). Furthermore, a number of GAox-related *2-*oxoglutarate-dependent dioxygenases (2-OGDs) were identified in *Physcomitrella patens* and *Selaginella moellendorffii* prompting the theory that non-canonical GA-oxidation reactions might catalyse the production of diverse GA-like compounds in bryophytes and tracheophytes (70, 91). The similarity in transcriptional responses between *M. paleacea* and *M. truncatula* and the induction of several GAox-related *2-OGDs* oxidases in our dataset encourages us to support the hypothesis that non-canonical GA-like diterpenoids are produced in bryophytes, and that the synthesis of these compounds is enhanced during symbiosis. As suggested by Cannel *et al.* (71), the lack of evidence for GA-like diterpenoids in bryophytes (69, 91) is limited to mosses and might be due to low levels of expression in the observed taxa, which are AMS non-hosts. As GA biosynthetic genes are significantly induced in AMS *M. paleacea*, an analysis of the diterpenoid content of colonized thalli might improve our understanding of the GA profile of early diverging land plants.

Whilst confirming the importance of several characterised molecular pathways, this study provides novel evidence to support several gene families with a predicted but yet uncharacterised mechanism of action in AMS: *OsNOPE1, MtRFCb, MtKIN3/OsARK1, MtKIN6, MtABCB20, MtDHY,MtGST1, MtDUF538, MtHYP5a, MtAP2/ERF2a*, *MtAmpB,* and several genes encoding cytochrome P450 family proteins and chitinases (22, 47, 92). These genes are strictly conserved across AMS-host angiosperms and transcriptionally induced during AMS in *M. truncatula,* some of them displaying abnormal mutant AMS phenotypes (22, 47, 92). Our study’s finding that these candidate AMS genes are both conserved and transcriptionally induced in *M. paleacea* supports the notion that they might be required for symbiosis and that they were recruited for AMS before the LCA of liverworts and angiosperms. Functional characterization of these candidates in *M. paleacea* might aid the discovery of their role in AMS, as the lower number of paralogs per orthogroup identified in *M. paleacea* might overcome issues of genetic redundancy previously observed in *M. truncatula* and other angiosperm models.

In conclusion, our study demonstrates that the genetic machinery regulating pre-symbiotic signalling, transcriptional reprogramming and nutrient exchange in arbuscular mycorrhiza symbiosis is largely conserved and coregulated across liverworts and angiosperms, despite more than 400 million years of divergence since the LCA of these species (93).

An increasing body of evidence suggests that bryophytes are a monophyletic clade, sister to all other land plants (23, 93–96). The implication of this evidence is that any genetic sequence evolutionarily conserved between an internal clade of bryophytes (e.g. liverworts) and an internal clade of tracheophytes (e.g. angiosperms), is ancestral to land plants. In the context of AMS, the evolutionarily conserved genes and pathways presented in this study are thus likely to be representative of the ancestral molecular toolkit that regulated AMS in the LCA of land plants. Going forward, comparative analyses of bryophyte and tracheophyte AMS models will allow us to pinpoint what aspects of AMS are ancestral to embryophytes, improving our understanding of the selective pressures and evolutionary landscapes that made AMS a selective advantage at the earliest stages of land plant evolution.

## Materials and Methods

### Plant materials and growth conditions

*Marchantia paleacea* wild-type thalli were provided by P. Carella and S. Schornack (Sainsbury Laboratory, University of Cambridge). Thalli and gemmae were grown axenically on half-strength Gamborg B5 medium (Duchefa Biocheme) in sterile 0.8% agar plates. Plants were grown in a controlled growth chamber at 22 °C, with a continuous light intensity of 100 μmol m−2 s−1 PAR (photosynthetically active radiation).

### AMS colonization assay

For AMS colonisation, 4-week old axenically grown *M. paleacea* thalli were transferred to (5×5×6 cm) pots containing sand and 5ml *R. irregularis* crude inoculum or 5ml of 2x autoclaved crude inoculum for mock-inoculated conditions. Inoculum was produced by prolonged co-culture of *Tagetes multiflora* and *R. irregularis* spores in sand (97). Inoculated and mock-inoculated plants were grown in a controlled growth chamber at 22 °C,16:8 h day-night cycles with a light intensity of 200 μmol m−2 s−1 PAR. Plants were watered three times per week with mistified “artificial rainwater solution” (pH 5.8) (98).

### Microscopy and fungal structure quantification

Thalli were collected at 5, 8 and 11 WPI for mock and *R. irregularis*-inoculated samples to monitor AMS colonization and stored in 50% ethanol. For Trypan blue staining, thalli were incubated overnight in 10% potassium hydroxide, washed 10x with distilled water, then incubated overnight in staining solution (50% lactic acid, 25% glycerol, 25% ddH20, 0.1% trypan blue). For sectioning, stained samples were incubated in destaining solution (50% lactic acid, 25% glycerol, 25% ddH20) for 1 hour then embedded in 3.5% agarose gel. Transversal sections (150-200μm) were taken at 10 equally distanced positions spanning the length of each thallus, using a Hyrax V50 vibratome (Zeiss, Oberkochen, Germany). Sections were imaged under a brightfield microscope. Fungal structures and cell-wall pigment accumulation in each biological replicate were quantified by assessing their presence or absence in each of the 10 transversal sections.

### RNA extraction and RNA-seq library preparation

Samples were collected at 5, 8 and 11 WPI, washed in distilled water and fixed in ice-cold 100% methanol. For each independent biological replicate four thalli were collected and pooled together for subsequent analysis. To maximise the ratio of colonized to non-colonized tissue in each sample, the midribs of collected thalli were excised to remove apical notches, lateral margins and gemma cups as these structures are not colonised by AMS fungi (51). Total RNA was extracted from the excised midribs using PureLink Plant RNA Reagent (Thermo Fisher Scientific, Waltham, USA) following manufacturer’s instructions. Addition of 2% Polyvinylpyrrolidone (PVP40) to the Plant RNA Reagent mixture before extraction improved RNA purity. RNA samples were treated with Turbo DNA-free DNase (Invitrogen-Thermo Fisher Scientific, Waltham, USA) to remove contaminating genomic DNA. cDNA library preparation was performed using 1μg of total RNA with a TruSeq Stranded mRNA Library Preparation Kit High Throughput (Illumina, San Diego, California, USA) according to manufacturer’s instructions (Catalog # RS-122-9004DOC, Part # 15031047 Rev.E). Library quality was assessed using a DNA1000 chip on Bioanalyzer 1200 (Agilent Technologies, Santa Clara, California, USA) and library quantities were measured with a Qubit dsDNA BR Assay Kit (Thermo Fisher Scientific, Waltham, USA). A total of 24 samples (*R. irregularis* and mock-inoculated conditions, three time points, 4 biological replicates) were pooled with 16 samples from a parallel experiment, multiplexed and sequenced on a NextSeq500 (Illumina, San Diego, California, USA) as a 2 × 75nt paired-end run (10 million reads per sample).

### Transcriptome assembly and annotation

Raw RNAseq reads were quality-filtered using FastQC v0.11.5 (https://github.com/s-andrews/FastQC) and adaptor sequences trimmed with Cutadapt v1.3 (99). For transcriptome assembly, all trimmed reads (400 million total) were mapped against the *M. paleacea* genome v1 (14) and the *R. irregularis* genome DAOM_181602_v1.0 (100) with HiSat2 v2.1.0 (101). The resulting SAM file was sorted and indexed using SAMtools v1.9 (102). Using this alignment, a reference transcriptome was assembled using Cufflinks v2.2.1 with multi-mapped read correction (103). The assembled transcriptome was annotated using Transdecoder v5.5.0 for protein-coding region prediction (104) and Trinotate v3.1.1 (55).

### Differential gene expression analysis

All sample libraries were mapped to the newly assembled *M. paleacea* and *R. irregularis* reference transcriptome with HiSat2 v2.1.0 (101). Reads were initially mapped to the reference transcriptome including both *M. paleacea* and *R. irregularis* transcripts, then reads mapping to *R. irregularis* scaffolds were filtered out of downstream analyses. Htseq-count (105) was used to quantify transcript levels using the newly assembled *M.paleacea* transcriptome as input annotation file. Differential expression analysis was performed with DESeq2 v1.22.2 (106), considering only genes with at least ten reads across all timepoints/conditions (26433 genes). Principal component analysis (PCA) was performed on the DESeq2 model to observe clustering of biological replicates, and one of five replicates for the ‘5W-MYC’ timepoint was identified as an outlier and removed from the analysis. Differentially expressed genes were identified by pairwise comparison of mock-inoculated vs. mycorrhizal samples at the same time-point, with a significance threshold of log2 fold-change > |1| and adjusted p-value < 0.05. Differentially expressed genes were used to perform hierarchical clustering of samples and plotted with the R pheatmap package (107) using variance-stabilised counts median-centered by gene.

### Gene ontology enrichment analysis

Gene set enrichment analysis was performed with OmixBox (108), using a two-tailed Fisher’s exact test with p-value correction by False Discovery Rate (FDR) control according to Benjamini-Hochberg (109). The significant threshold for GO enrichment was set as FDR < 0.05.

### Orthology inference and comparative analysis

Gene orthology inference was performed with OrthoFinder-v2.3.3 (110), using the *M. paleacea* transcriptome assembled in this study and 21 publicly available plant protein datasets (Supplementary table 3). The orthogroups and the orthologs identified through OrthoFinder were then used to assess evolutionary conservation of *M. truncatula* orthologs in *M. paleacea*. For this analysis we selected the time point with the strongest response to AMS in our RNAseq dataset (11WPI) and compared it to the timepoint with the strongest response to AMS in Luginbuehl *et al*.’s RNAseq dataset (27DPI)(19). We considered the colonization stages in the different species to be comparable as they displayed all fungal structures associated with late symbiosis. After differential expression analysis, we first compared the Log2 FC of SCOs DE (P-adj < 0.05) in both datasets (Fig. 2a). We subsequently expanded our analysis to include orthologs with one copy in *M. paleacea* and multiple copies in *M. truncatula,* subsetting the dataset to include only DE genes in *M. paleacea* with at least one DE ortholog in *M. truncatula* (Supplementary table 6). The phylogenetic tree of *MPA7307* was constructed using all protein sequences within the *MPA7307* orthogroup (OG0000006). Amino acid sequences were aligned with MAFFT v7 (111) and maximum-likelihood tree inference was performed with IQ-TREE 2 (112) using the JTT + I + G4 amino acid substitution model. The best-fitted evolutionary model was identified using ModelTest-NG (113). Bootstrapping was performed with 1000 replicates of Ultrafast Bootstraps (114) and 1000 replicates of SH-like approximate likelihood ratio test (115).

## Acknowledgments

We thank Sebastian Schornack and Philip Carella for the *M. paleacea* plant material and for support with establishing *M. paleacea* cultures and techniques, Giles Oldroyd and Guru Radhakrishnan for kindly sharing *M. paleacea* genomic resources, and Hajk-Georg Drost for guidance with comparative analysis. M.S. was supported by the UK Biotechnology and Biological Sciences Research Council (BBSRC) (BB/M011194/1). U.P. is supported by the research project Engineering the Nitrogen Symbiosis for Africa (ENSA), which is funded by a grant to the University of Cambridge by the Bill & Melinda Gates Foundation and the Foreign, Commonwealth & Development Office (FCDO).

## Data availability

Transcriptome sequences (nucleotides and predicted peptides) assembled and discussed in this work are provided in Supplementary Data. The resolved gene trees for each orthogroup presented in Supplementary Table 4 are included in Supplementary Data. The raw data from transcriptomics is being deposited on GEO.

