## Supplementary Figures for "Transcriptional responses to arbuscular mycorrhizal symbiosis development are conserved in the early divergent *Marchantia paleacea*"

### Supplementary information

### Supplementary figures

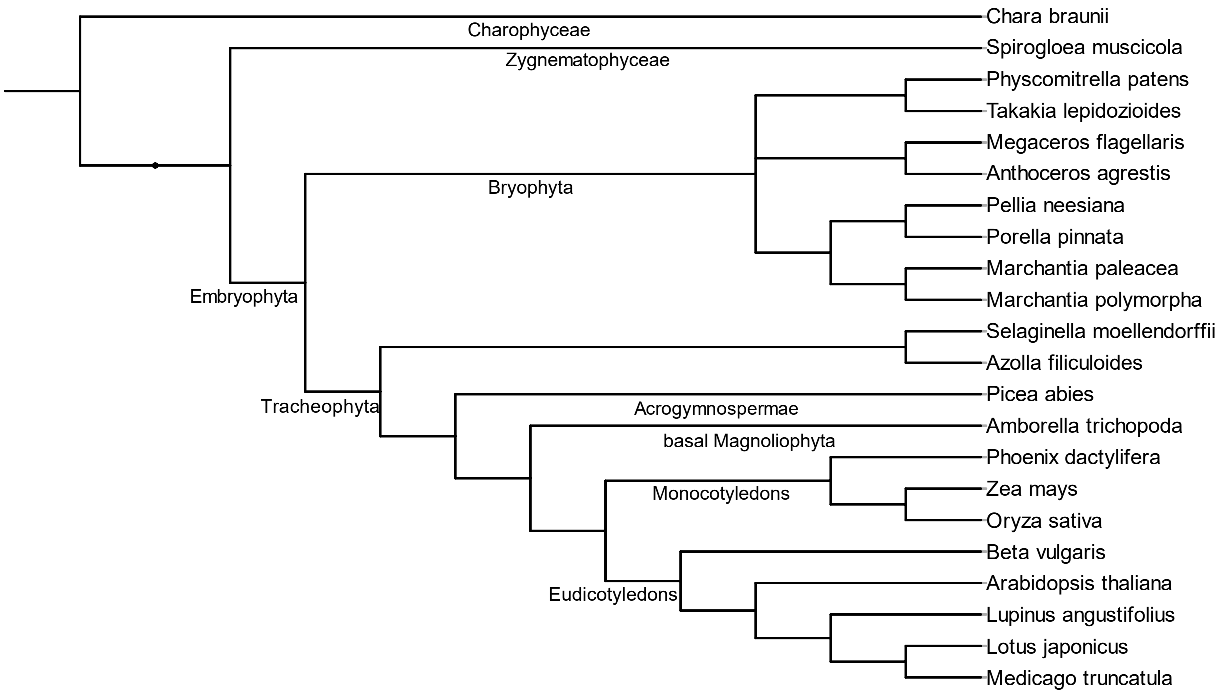


**Fig. S1. Cladogram of plant species used for Orthofinder2 phylogenetic analysis.** Major taxonomic groups are indicated on the branches.

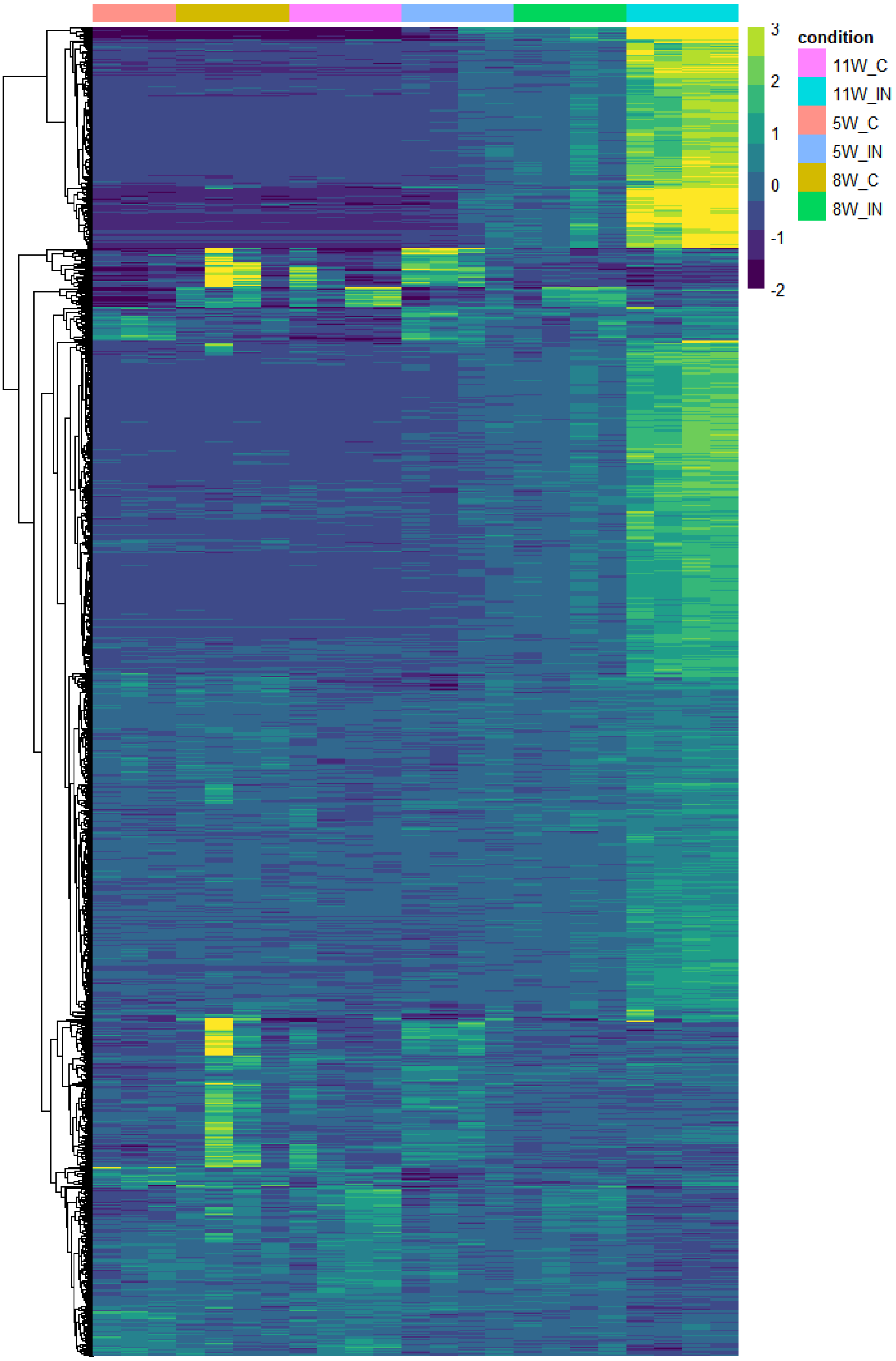


**Fig. S2. Hierarchical clustering of differentially expressed *M. paleacea* genes (L2FC>|1|, adjusted P-value < 0.05) with a corresponding *M. truncatula* ortholog upregulated during AMS colonization (1).** Variance-stabilised counts median-centered by gene are shown**.**

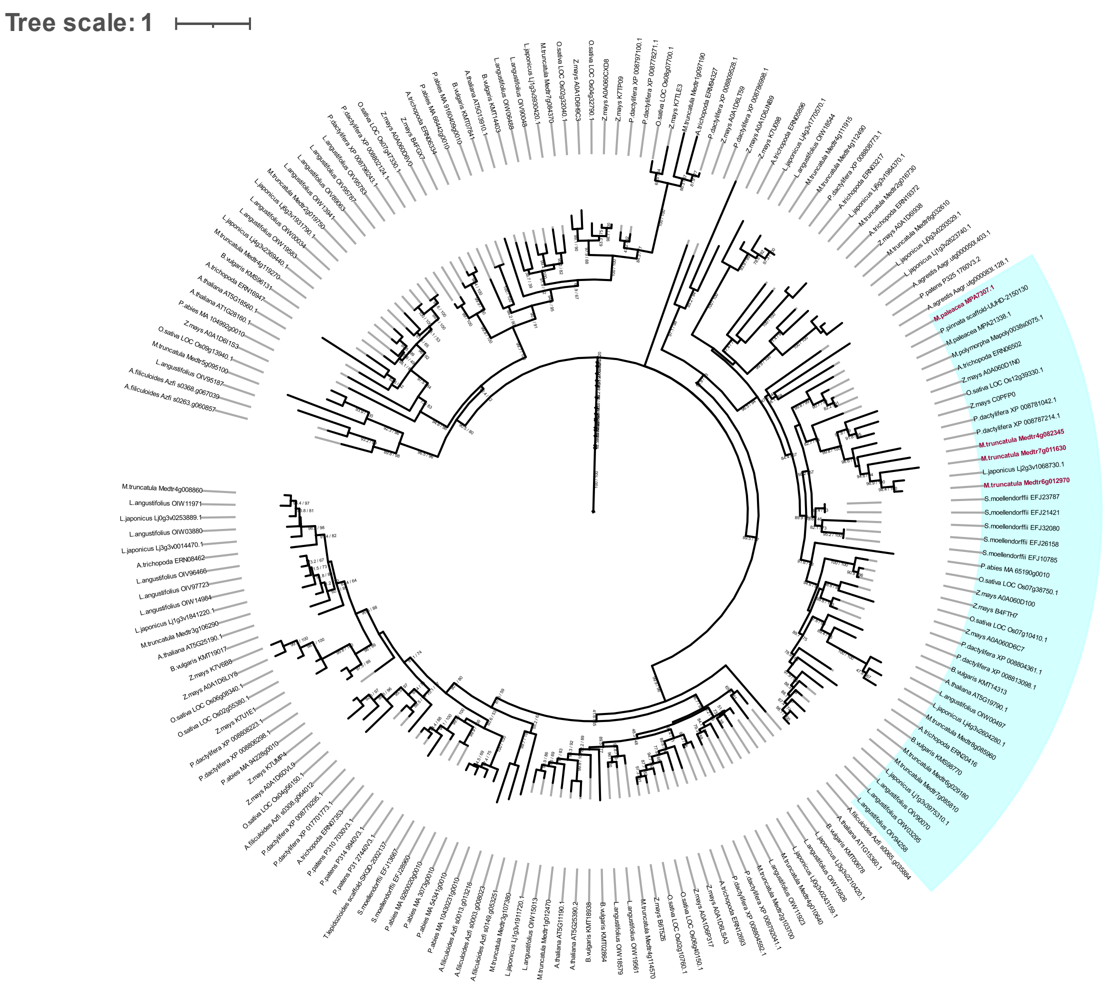


**Fig. S3. Maximum likelihood phylogenetic tree of the RAP211 family of AP2/ERF domain transcription factors.** The phylogenetic tree sown is a clade from the bigger AP2/ERF superfamily phylogenetic tree (orthogroup: OG0000006). Branch support from Ultrafast Bootstraps (dividend) and SH-like approximate likelihood ratio test (divisor) is shown. The *M. paleacea* and *M. truncatula* orthologs of genes lost from the genomes of AMS non-host species are highlighted in magenta.
